# Four-dimensional characterization of the *Babesia divergens* asexual life cycle: from the trophozoite to the multiparasite stage

**DOI:** 10.1101/2020.07.23.219030

**Authors:** José Javier Conesa, Elena Sevilla, María C. Terrón, Luis Miguel González, Jeremy Gray, Ana J. Pérez-Berná, José L. Carrascosa, Eva Pereiro, Francisco Javier Chichón, Daniel Luque, Estrella Montero

**Author notes:** Address correspondence to Daniel Luque and Estrella Montero. José Javier Conesa and Elena Sevilla contributed equally to this work. Author order was determined on the basis of seniority.

## Abstract

*Babesia* is an apicomplexan parasite of significance that causes the disease known as babesiosis in domestic and wild animals and in humans worldwide. *Babesia* infects vertebrate hosts and reproduces asexually by a form of binary fission within erythrocytes/red blood cells (RBCs), yielding a complex pleomorphic population of intraerythrocytic parasites. Seven of them, clearly visible in human RBCs infected with *Babesia divergens*, are considered the main forms and named single, double and quadruple trophozoites, paired and double paired-pyriforms, tetrad or Maltese Cross, and multiparasite stage. However, these main intraerythrocytic forms coexist with RBCs infected with transient parasite combinations of unclear origin and development. In fact, little is understood about how *Babesia* builds this complex population during its asexual life cycle. By combining the emerging technique cryo soft X-ray tomography and video microscopy, main and transitory parasites were characterized in a native whole cellular context and at nanometric resolution. As a result, the architecture and kinetic of the parasite population has been elucidated. Importantly, the process of multiplication by binary fission, involving budding, was visualized in live parasites for the first time, revealing that fundamental changes in cell shape and continuous rounds of multiplication occur as the parasites go through their asexual multiplication cycle. Based on these observations, a four-dimensional (4D) asexual life cycle model has been designed highlighting the origin of the tetrad, double paired-pyriform and multiparasite stages and the transient morphological forms that, surprisingly, intersperse in a chronological order between one main stage and the next along the cycle.

**IMPORTANCE:** Babesiosis is a disease caused by intraerythrocytic Babesia parasites, which possess many clinical features that are similar to those of malaria. This worldwide disease, is increasing in frequency and geographical range, and has a significant impact on human and animal health. *Babesia divergens* is one of the species responsible for human and cattle babesiosis causing death unless treated promptly. When *B. divergens* infects its vertebrate hosts it reproduces asexually within red blood cells. During its asexual life cycle, *B. divergens* builds a population of numerous intraerythrocytic (IE) parasites of difficult interpretation. This complex population is largely unexplored, and we have therefore combined three and four dimensional (3D and 4D) imaging techniques to elucidate the origin, architecture, and kinetic of IE parasites. Unveil the nature of these parasites have provided a vision of the *B. divergens* asexual cycle in unprecedented detail and a key step to develop control strategies against babesiosis

## INTRODUCTION

Babesia is an apicomplexan parasite which infect the red blood cells (RBCs) of a wide range of vertebrates, causing babesiosis (1). This disease, transmitted by ticks, has a significant impact on human and animal health. Two billion cattle worldwide are exposed to the infection causing substantial economic losses. The disease is also an emergent zoonosis of humans (2–4). *Babesia divergens*, is the most important species in Europe, causing redwater fever in cattle and severe and often fatal babesiosis in humans (1).

Once the vertebrate host has been bitten by an infected tick, sporozoites invade RBCs and begin an asexual life cycle known as merogony. This cycle has been partially elucidated and involves RBC invasion, metabolism and replication by a form of binary fission involving budding, resulting in merozoites that egress and destroy the host cell to seek and invade new uninfected RBCs (uRBCs) within seconds to minutes, thus perpetuating the infection (5, 6).

After several rounds of replication, *B. divergens* builds a complex population of distinct morphological intraerythrocytic (IE) stages. Seven of them, clearly distinguishable in the peripheral blood of infected humans and under *in vitro* growth conditions, are considered the main IE stages (7), namely single round trophozoite, paired-pyriforms (two attached pear-shaped sister cells), double trophozoites (two round unattached cells), double paired-pyriforms (two sets of paired sister cells), tetrads or Maltese Crosses (four attached sister cells), quadruple trophozoites (four round unattached cells) and multiple parasites (RBCs containing more than four parasites).

The merogony of *Babesia* is asynchronous and IE parasites in different stages coexist with free merozoites in the bloodstream (3). Despite the asynchronous nature of *B. divergens* replication, an approximation to the putative morphogenetic pathway was derived from *in vitro* methods alongside visible light microscopy, which allowed the visualization of a synchronized *B. divergens* asexual cycle for the first 24 hours, but exclusively involving the seven IE main stages (7). Sequentially, after the invasion of RBCs by free merozoites, the resulting single trophozoites give rise to paired-pyriforms. Then, paired-pyriforms develop into double trophozoites, which may give rise to double paired-pyriforms. Paired-pyriforms occasionally differentiate to tetrads or Maltese Crosses. Double paired-pyriforms and tetrads result in quadruple trophozoites and finally, quadruple trophozoites differentiate into multiple parasites. However, 24 hours later, the life cycle progresses and lose its synchronicity, transforming into asynchronous populations in highly parasitized RBCs that contain transient morphological parasites of unclear origin alongside the seven main IE stages, mimicking the situation in human infections.

In spite of recent advances, our comprehension of the asexual *B. divergens* life cycle, including the multiplication process that a single parasite undergoes within an original iRBC to develop ultimately into a multiparasite stage or the origin of tetrads, is still hampered by the limited knowledge of the kinetics and morphology of the parasite, currently based on information from light and electron microscopy of fixed and stained ultrathin sections (6–10). In this context, cryo-soft X-ray tomography (cryo-SXT) is a tool that bridges the gap between light and electron microscopy resolving some challenges in imaging and making unnecessary the use of contrasting agents, thus avoiding sectioning and staining artefacts (11–13). Taking into account these advantages, here we use cryo-SXT to obtain the three-dimensional (3D) reconstructions of cryo-preserved, intact (non-sectioned) unstained *B. divergens* infected RBCs (iRBCs) under close-to-native-state conditions. These data reveal not only the 3D architecture of the known main seven IE stages in their native environment, but also novel transient IE parasites from *B. divergens* asynchronous cultures in a whole cell context, and at nanometric resolution.

In addition to a detailed morphological description of *B. divergens* using cryo-SXT, our study is complemented by video microscopy over time (4D imaging) and transmission electron microscopy (TEM), thus providing an insight in the kinetics to reveal a comprehensive model of the complex *B. divergens* asexual cycle that differ in some aspects to the previously proposed models (7, 14, 15). During this dynamic and pleomorphic *in vitro* cycle, it is possible to observe how *B. divergens* induces several cytological events that explain the origin and development of the main IE stages as well as the role of the transient morphological parasites that surprisingly intersperse between one main IE stage and the next one in the cycle.

## RESULTS

### Three-dimensional (3D) structure of the *B. divergens* blood stages

To characterize the 3D architecture of *B. divergens*, stained fluorescent parasites from in *vitro* asynchronous cultures were analyzed by correlative visible light fluorescence microscopy and cryo-SXT (Fig. 1). Acquisition of more than 200 cryo-SXT datasets were required to deal with the variety and complexity of the *B. divergens* pleomorphic forms involved throughout the parasite asexual life cycle.

**Fig. 1.**
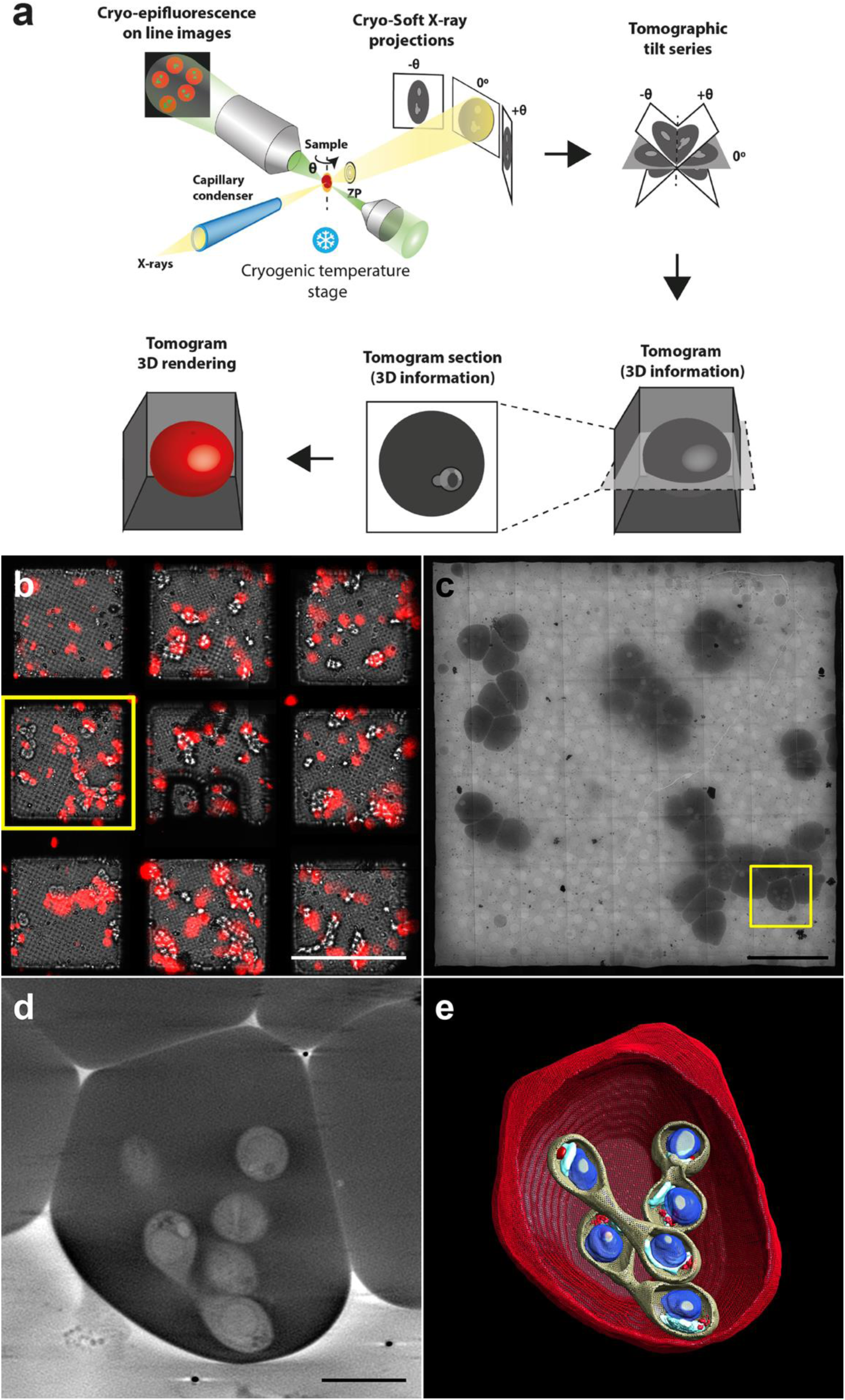
Correlative cryo-epifluorescence and cryo soft X-ray tomography imaging of *B. divergens* human iRBCs. (a) This correlative workflow, used at MISTRAL beamline at ALBA synchrotron, provides biological images and structural information of whole *B. divergens* human iRBCs close to their native state at a spatial resolution of around 50 nm. *B. divergens* iRBCs are tilted to different angles and an image is acquired at each angle. The tilt-series of images are reconstructed into a three-dimensional (3D) tomogram providing structural information of the whole cells. (b) *B. divergens* asynchronous *in vitro* cultures labelled with MitoTracker red (red fluorescence) are deposited on to holey carbon EM grids in an optimal cell confluency (10^5^ cells per grid) and plunge-frozen in liquid ethane. The vitrified grids are screened with an online epifluorescence microscope to generate a fluorescence map and select the most relevant cells (yellow square). (c) Grids are loaded into the MISTRAL transmission X-ray microscope at the ALBA synchrotron light source for screening. An X-ray mosaic of projection images is generated and the previous fluorescence map helps in locating again the same cells in the yellow square. (d) Cryo-SXT tomogram sections of a *B. divergens* multiparasite stage acquired in the yellow squared area in c. (e) 3D rendering of the acquired cryo-SXT tomogram shown in d. The scale bars of b, c and d are 100 µm, 20 µm and 2 µm, respectively.

Reconstructed tomograms were used to recover the intracellular 3D cartography of the main seven IE stages (Fig. 2 and Fig. 1). Three-dimensional data showed a morphological shape characteristic of each parasite stage. Thus, free merozoites are polarized ellipsoidal cells with an apical width prominence at the end (Fig. 2a-b). Single (Fig. 2c-d), double (Fig. 2e-f) and quadruple trophozoites (Fig. 2g-h) showed a round shape, while paired-pyriforms (Fig. 2i-j), tetrads (Fig. 2k-l) and double paired-pyriforms (Fig. 2m-p) exhibited the characteristic pear-shaped form. Some 3D sub-cellular compartments were clearly discernible in both free merozoites and IE stages including an elongated mitochondrion (1-2 µm) and a round apicoplast (300 nm size) next to the nucleus (700 nm size), which occupied most of the parasite cytoplasm. Dense granules were positioned on one side of the round trophozoites or close to the apical end of free merozoites and pear-shaped parasites (Fig. 2). It was not possible to resolve clearly the 3D structure of the Golgi apparatus and the endoplasmic reticulum (ER) due to the resolution attained (around 50 nm in 3D). We also detected by cryo-SXT other intracellular structures in free merozoites and IE stages that may correspond to micronemes, rhoptries and the inner membrane complex (IMC), previously observed by TEM (8, 9). However, it was not possible to unequivocally identify and/or count them (Fig. S1).

**Fig. 2.**
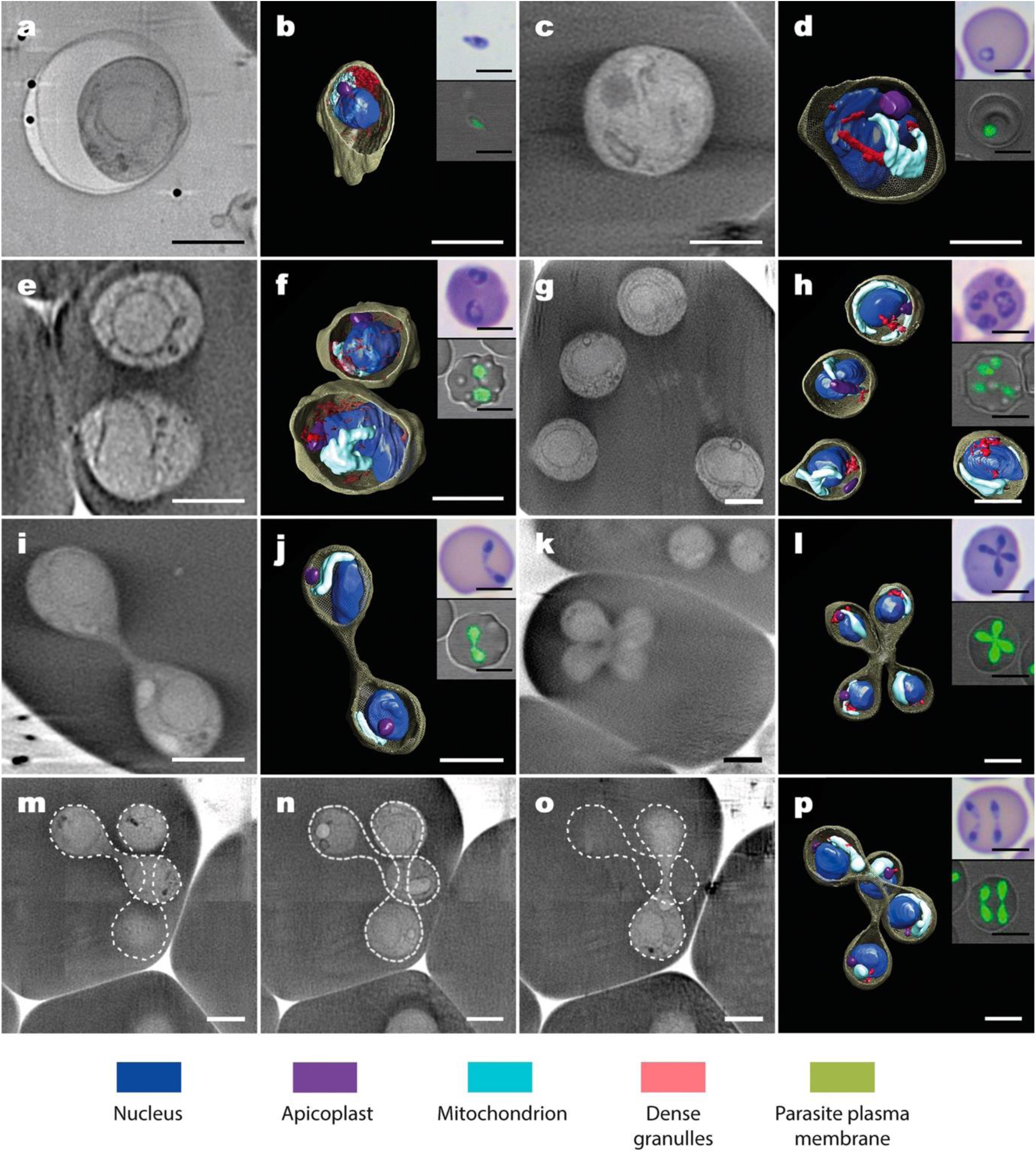
Three-dimensional (3D) architecture of intact *B. divergens* parasites. Panels show the cryo-SXT tomogram sections of the *B. divergens* free merozoite and six main stages within human RBCs. The corresponding 3D tomogram renderings show the architecture of intact parasites and the organelle distribution in colour. Insets show the equivalent parasite stages stained with Giemsa and observed by light microscopy (top inset) or stained with MitoTracker (green fluorescence) and observed using a confocal laser microscope (lower inset). (a, b) Free merozoite. (c, d) Single round trophozoite stage. (e, f) Double round trophozoite stage. (g, h) Quadruple round trophozoite stage. (i, j) Paired-pyriform stage. (k-l) Tetrad (Maltese Cross). (m, n, o) Tomogram sections of a double paired-pyriform stage at different depths over the same 3D reconstruction. (p) 3D tomogram rendering of the corresponding double paired-pyriform stage. The scale bars of panels a-p are 2 µm. The scale bars of insets from panels a-p are 5 µm.

Other membranous systems were clearly visible in the iRBCs. These include i) possible hemoglobin-containing vesicles within the parasite cytoplasm and ii) low absorbing vesicles with sub-micron size and novel long membrane structures, both within the cytoplasm of iRBCs. Thus, a single round dense feature was detected exclusively in the cytoplasm of trophozoites (Fig. S2a-b). Since these dense structures exhibited a similar X-ray linear absorption coefficient to the one of hemoglobin from the RBC cytoplasm, we hypothesized that they could be hemoglobin-containing vesicles. It is interesting to note that similar hemoglobin inclusions, and the possible parasite endocytic uptake of the hemoglobin from the cytoplasm of iRBCs, were observed in TEM serial sections (Fig. S2c-h). These membranous structures were heterogeneous in size (250-600 nm) and could be the result of the internalization of an RBC cytoplasm portion to form the hemoglobin-containing vesicle.

The sub-micron vesicles present in the cytoplasm of iRBCs showed different sizes (120, 250 and 400 nm), and some of them were also observable by Cryo-SXT and TEM (Fig. S3). Long structures (1.5-eared as a unique membranous extension. This feature extended from the parasite plasma membrane to the RBC plasma membrane, establishing a connection between the parasite and RBC (Fig. S4).

In addition to finding and recognizing the seven main IE stages in a whole cell context, we observed other novel IE transient morphological forms with a complex pleomorphic 3D architecture. The elucidation of the origin and role of these new IE forms in the parasite life cycle was addressed using a combination of cryo-SXT and long-term time-lapse video microscopy, as described below.

### Intraerythrocytic asexual cycle of *B. divergens*: from the trophozoite to the paired-pyriform

Both asynchronous *B. divergens in vitro* culture and in peripheral blood of humans reflect a confused scenario of a heterogeneous parasite population when seen by standard microscopic techniques (Giemsa stain and light microscopy, and TEM). To define a comprehensive and chronological organization of these IE forms in the cycle, beginning with the single trophozoite development after RBC invasion and ultimately ending with the multiparasite stage formation, we filmed the asynchronous *B. divergens* culture for long periods and combined video microscopy and cryo-SXT data.

We captured images of newly iRBCs and RBCs already parasitized with single trophozoites. Video microscopy showed how these single trophozoites reproduced by a form of transverse binary fission that involves budding. Some details were also observable by cryo-SXT. In a first phase of development, trophozoites adopted amoeboid shapes (Fig. 3a and 2g) until they reached a round form with two protuberant buds (Fig. 3b and 2h). This form was previously observed by TEM in *B. divergens* and more recently in *B. bigemina* and was named budding form (“Mickey Mouse”) because the buds contained organelles, indicating the posterior development of potential merozoites (9, 10). Video microscopy showed, step by step, how the budding form preceded the ultimate paired-pyriform development in a second phase. During this second phase, the budding form underwent a large change in morphology and became elongated; meanwhile, a transverse constriction was formed around the middle of the body (Fig. 2i and 3c). The initial transverse constriction progressively changed to a protuberant knob that ultimately developed a narrow waist (Fig. 2j, 3d, and S5). This fine structure divided the main body into two attached pear-shaped sister cells of equal size forming the paired-pyriform stage (Movie S1 at https://figshare.com/s/8ba6afd9e161899d682c). According to the cryo-SXT and TEM data, there are organelles and subcellular structures located in the zone that connect both sister cells. It seems that the development of the transverse constriction (Fig. 2i and 3c) and the distribution of cell material contained inside (Fig. 3n and S5) occurred as a coordinated process resulting in the two identical attached cells, each of them with a complete set of organelles at the end of the process (Fig. 3o). Cryo-SXT 3D reconstructions also allowed us to visualize how trophozoite cartography changes in order to yield a paired-pyriform (Fig. 3l-o). Notably, after imaging the morphogenesis of the trophozoite during its development by video microscopy and cryo-SXT, we were able to correctly identify and sequentially organize the corresponding IE forms when we saw them by light microscopy (Fig. 3q-t).

**Fig. 3.**
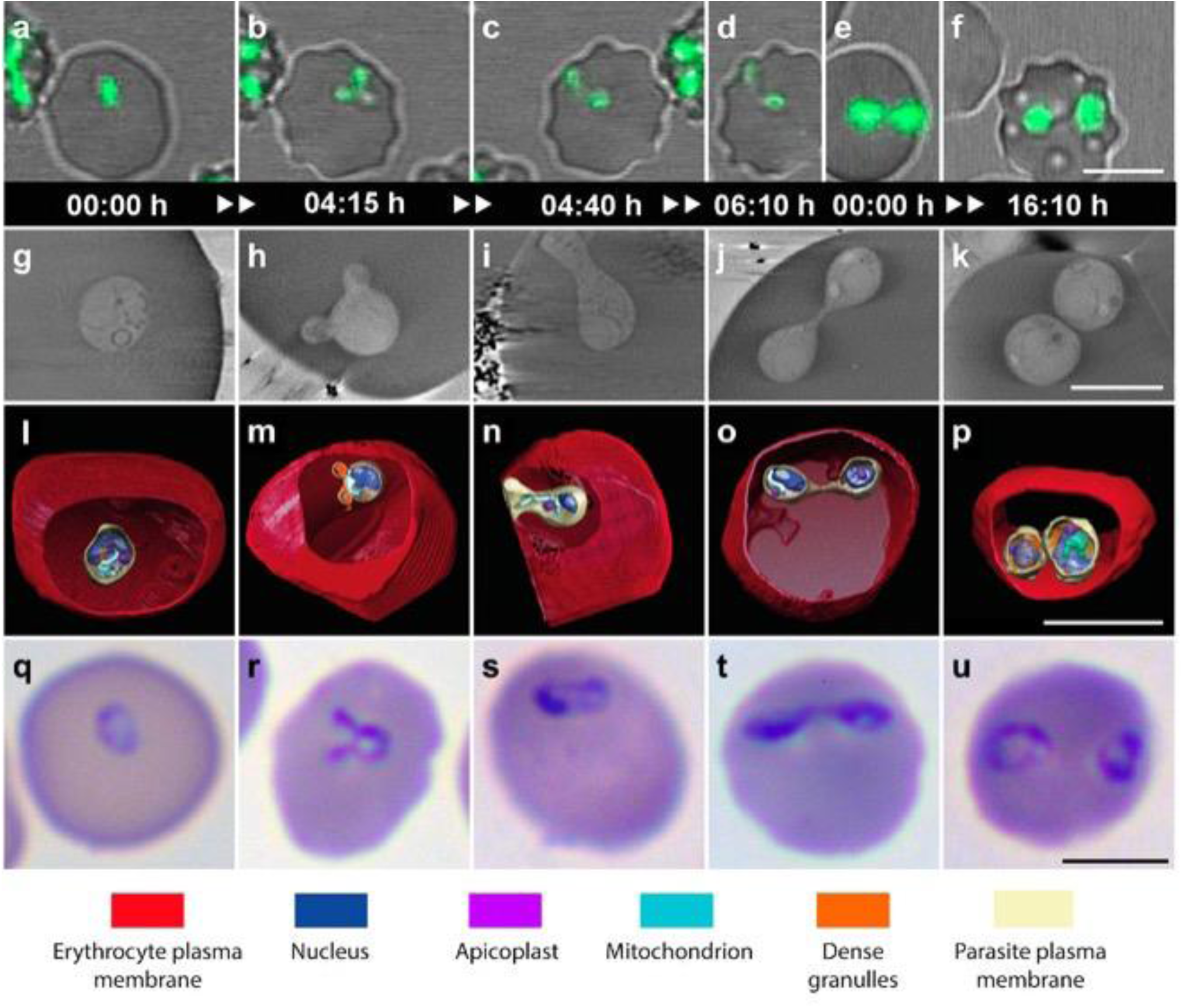
Dynamic development of single trophozoite and paired-pyriform stages. The figure shows the identification and development of the single trophozoite and paired-pyriform stages and the transient forms that intersperse in a chronological order between both main stages within the human RBC. (a-f) Time-lapse image sequences, captured by video microscopy, of *B. divergens* parasites stained with MitoTracker (green fluorescence) within the human RBC. Equivalent IE parasite forms are identified in *in vitro* cultures by cryo-SXT and Giemsa staining and light microscopy. (g-k) Cryo-SXT tomogram sections of the *B. divergens* iRBCs. (l-p) The corresponding 3D tomogram renderings show the architecture of intact parasites and the organelle distribution in colour. (q-u) *B. divergens* iRBCs stained with Giemsa. Panels are organized sequentially according to the video microscopy data. (a, b) The single trophozoite adopts amoeboid shapes until reaches the budding form at 4h and 15 min. (c) The budding form develops into an early paired pyriform at 4h and 40 min. (d) The early paired-pyriform ultimately develops into a paired-pyriform stage at 6h and 10 min. (e, f) The paired-pyriform stage develops into the double trophozoite stage. (g, l, q) Single trophozoite stage. (h, m, r). Budding form. (i, n, s) Early paired-pyriform under development. (j, o, t) Paired-pyriform stage. (k, p, u) Double trophozoite stage. (h-m) Budding form showing the initial segregation of cell material that appears concentrate in both buds. (n) Detail of the transverse constriction of the main cellular body and the presence of some organelles across the constriction zone. (o) Each daughter pear-shaped cell inherits a complete set of organelles at the end of the binary fission. The scale bars of a-k and q-u are 5 µm. The scale bars of l-p are 2 µm. Time-lapse imaging was captured every 5 min. The time lapse between each frame is indicated in hours and minutes (see also Movie S1 at https://figshare.com/s/8ba6afd9e161899d682c).

Moreover, by combining video microscopy and cryo-SXT, we have obtained the first 4D model description of the *B. divergens* asexual cycle starting with invasion by the free merozoite (Fig. 4a) followed by development of the single trophozoite into a paired-pyriform within the human RBC (Fig. 4b-f and j).

**Fig. 4.**
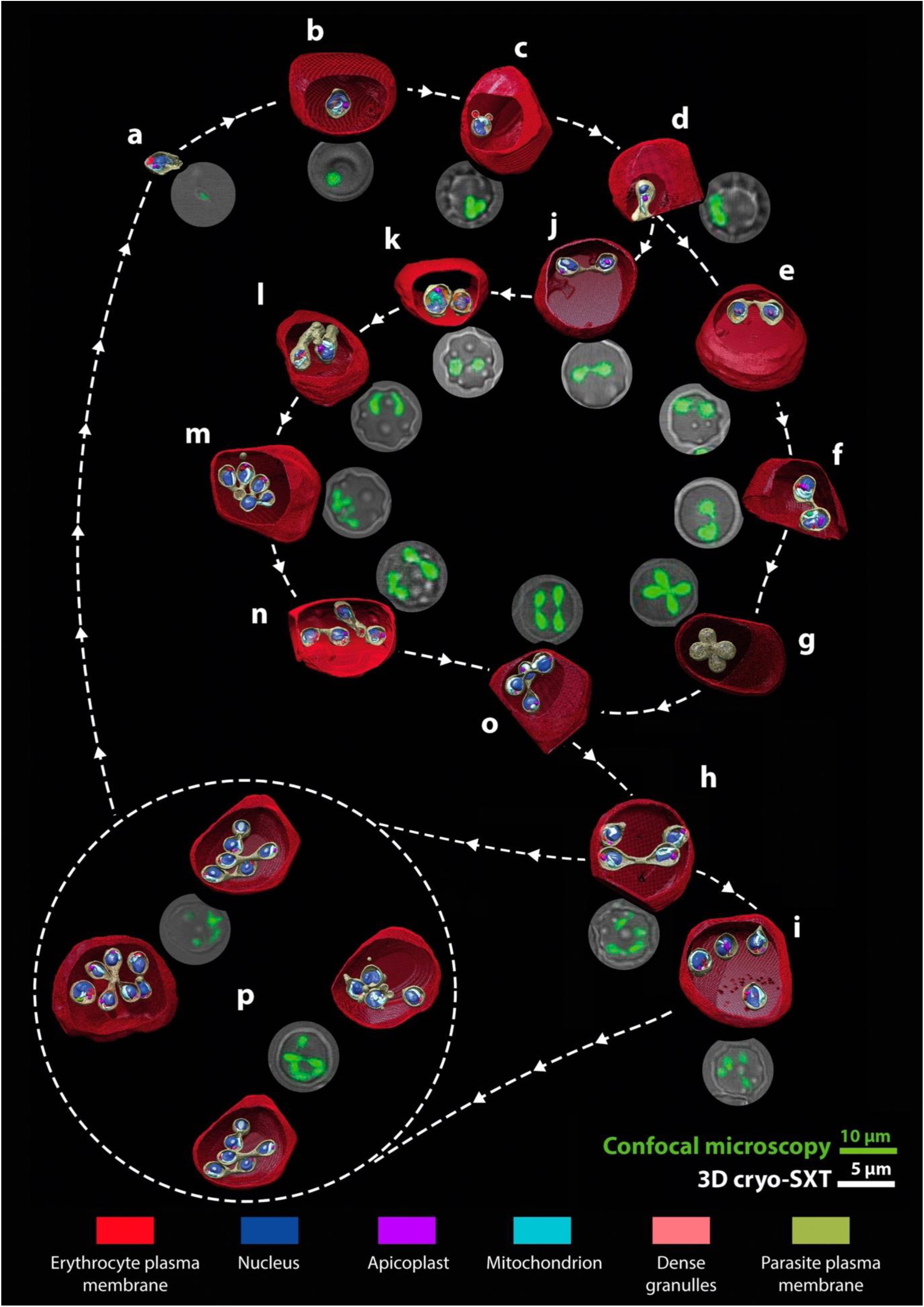
Four dimensional (4D) reconstruction of the main and intermediate intraerythrocytic (IE) stages that encompass the *Babesia divergens* asexual cycle. The cycle model shows a detailed chronological development that main IE stages undergo within the human RBC after the free merozoite invasion. (a-p) 3D rendering of cryo-SXT tomograms are organized according to the time-lapse images generated for parasites at different stage of development (green fluorescence). Cellular compartments and the distribution of the parasite organelles are indicated in colour. (a) A free merozoite about to invade a new human RBC. (b) Single trophozoite (main stage). (c) Budding form. (d) Early paired-pyriform under development. (e) Paired-pyriform (main stage). (f) Initial development process of a paired-pyriform into a tetrad. (g) Tetrad (main stage). (h) Two unattached pear-shaped parasites and a paired-pyriform. (i) Quadruple trophozoites (main stage). (j) Paired-pyriform (main stage). (k) Double trophozoites (main stage). (l) Two trophozoites adopting amoeboid forms during their development into double paired-pyriforms. (m) Double budding form. (n, o) Double paired-pyriforms (main stage). (p) Multiparasite stages (main stage). IE: intraerythrocytic.

### The paired-pyriform dominates the asexual cycle of *B. divergens*

After elucidating the dynamic development of the trophozoite, we continued analyzing the asexual cycle using the same experimental approach and following the chronological order established by (7). Hence, we captured images of RBCs parasitized with paired-pyriforms entering into a dynamic cycle of multiple pathways. Instead of egressing from the host cell (6), an alternative development of the paired-pyriform was to divide transversely yielding two pear-shaped cells. Several hours later, both cells adopted the characteristic round shape of the double trophozoite stage. (Fig. 3e-f, j-k, o-p, t-u, Fig. 4k-j, Fig. S6a and Movie S2 at https://figshare.com/s/8ba6afd9e161899d682c). Our recent studies showed that the dissociation of single paired-pyriforms into two pear-shaped sister cells occurs along the fine waist in a few seconds (6). However, the biomechanical process is not completely characterized and needs further study to understand the separation process.

Continuing with the cycle reconstruction, double trophozoites were transformed into double paired-pyriforms (Fig. 4k-o). During this dynamic process, each trophozoite multiplied by binary fission following the morphogenesis described above for a single trophozoite, but not necessarily simultaneously (Fig. S6b and S7, Movies S3 and S4 at https://figshare.com/s/8ba6afd9e161899d682c).

Of considerable interest was the development of paired-pyriforms into tetrads. Instead of dividing transversely, both sister pear-shaped cells remained attached and multiplied simultaneously yielding an attached double budding form or double “Mickey Mouse” that ultimately developed into a tetrad (Fig. 4e-g, S8a and Movie S5 at https://figshare.com/s/8ba6afd9e161899d682c). We also observed that tetrads can separate, yielding double paired pyriforms (Fig. 4g, 4o, S8b and S9a; Movies S6 and S7 at https://figshare.com/s/8ba6afd9e161899d682c) but we did not detect tetrads developing into quadruple trophozoites as previously suggested (7).

Less frequently, the development of double paired-pyriforms was observed to develop into quadruple trophozoites (Fig. 4o, 4h-i, Movie S8 at https://figshare.com/s/8ba6afd9e161899d682c).

### Double paired-pyriforms and quadruple trophozoites are involved in the development of multiparasite *B. divergens* stages

After formation of double paired-pyriforms and quadruple trophozoites, the cycle continued with the development of multiparasite stages (Fig. 4h and S9b; Movie S9 at https://figshare.com/s/8ba6afd9e161899d682c). Double paired-pyriforms and quadruple trophozoites underwent consecutive rounds of multiplication following a complex pathway of development and resulting in different multiparasite stages or polyparasitized RBCs infected with multiple combinations of parasite forms (Fig. 4p, and S10a; Movie S10) at https://figshare.com/s/8ba6afd9e161899d682c. Notably, multiparasite stages underwent new rounds of multiplication before ultimately egressing from the host cell (Fig. S10b, Movie S11 at https://figshare.com/s/8ba6afd9e161899d682c).

Finally, the time IE parasites took to transform into the next stages was measured and evaluated (Table 1). Of special interest is the finding that, regardless of the stage and the asynchronous multiplication of two or more parasites within the same RBC, the time elapsed from the budding form to the resulting new stage was similar. That is, trophozoites from any stage took similar times from the budding form to the paired-pyriform (1 h 38 min ± 48 min). This time was also comparable to the time required for the paired-pyriforms when developed through a synchronized budding to tetrads (Table 1).

**Table 1.**
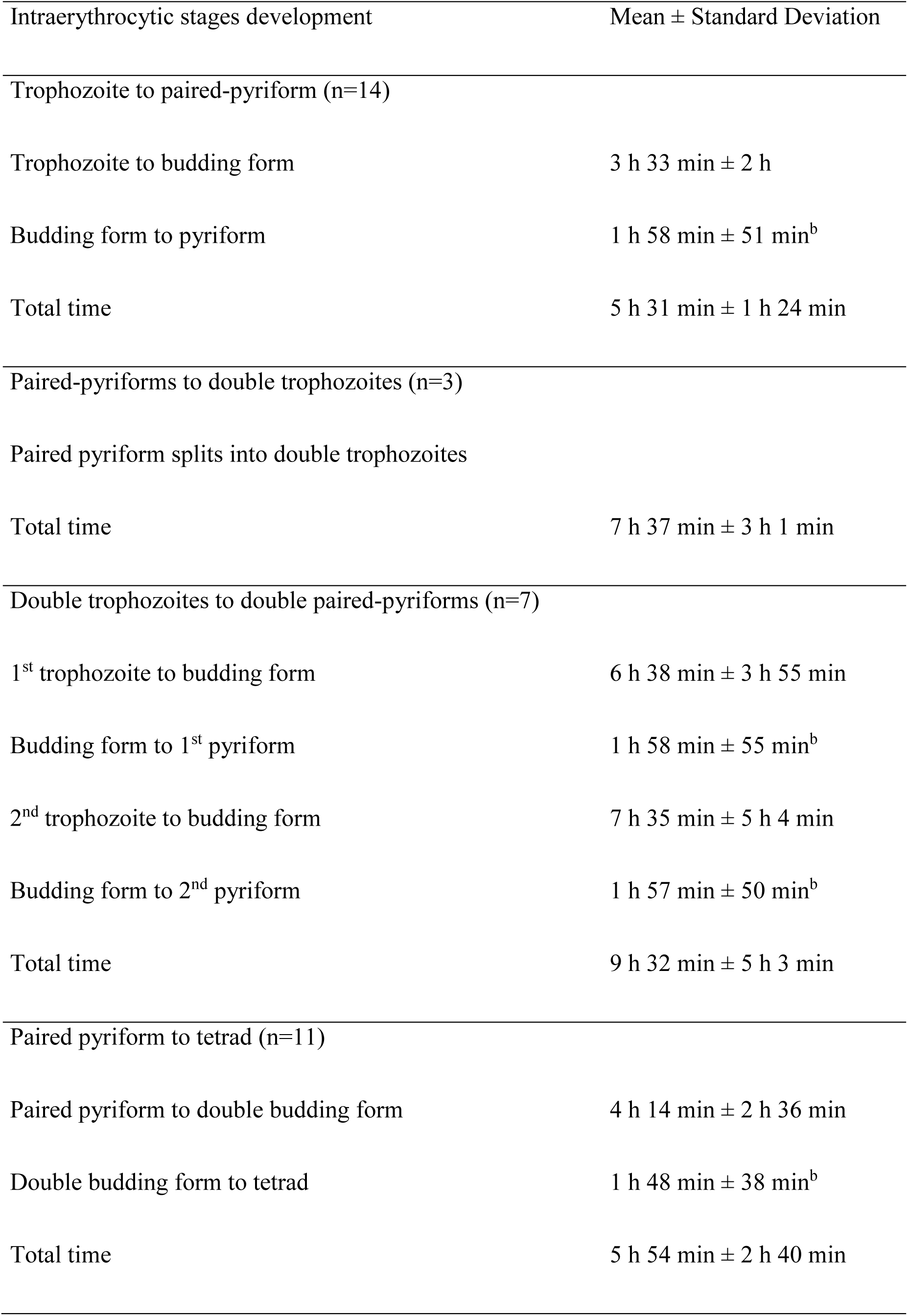
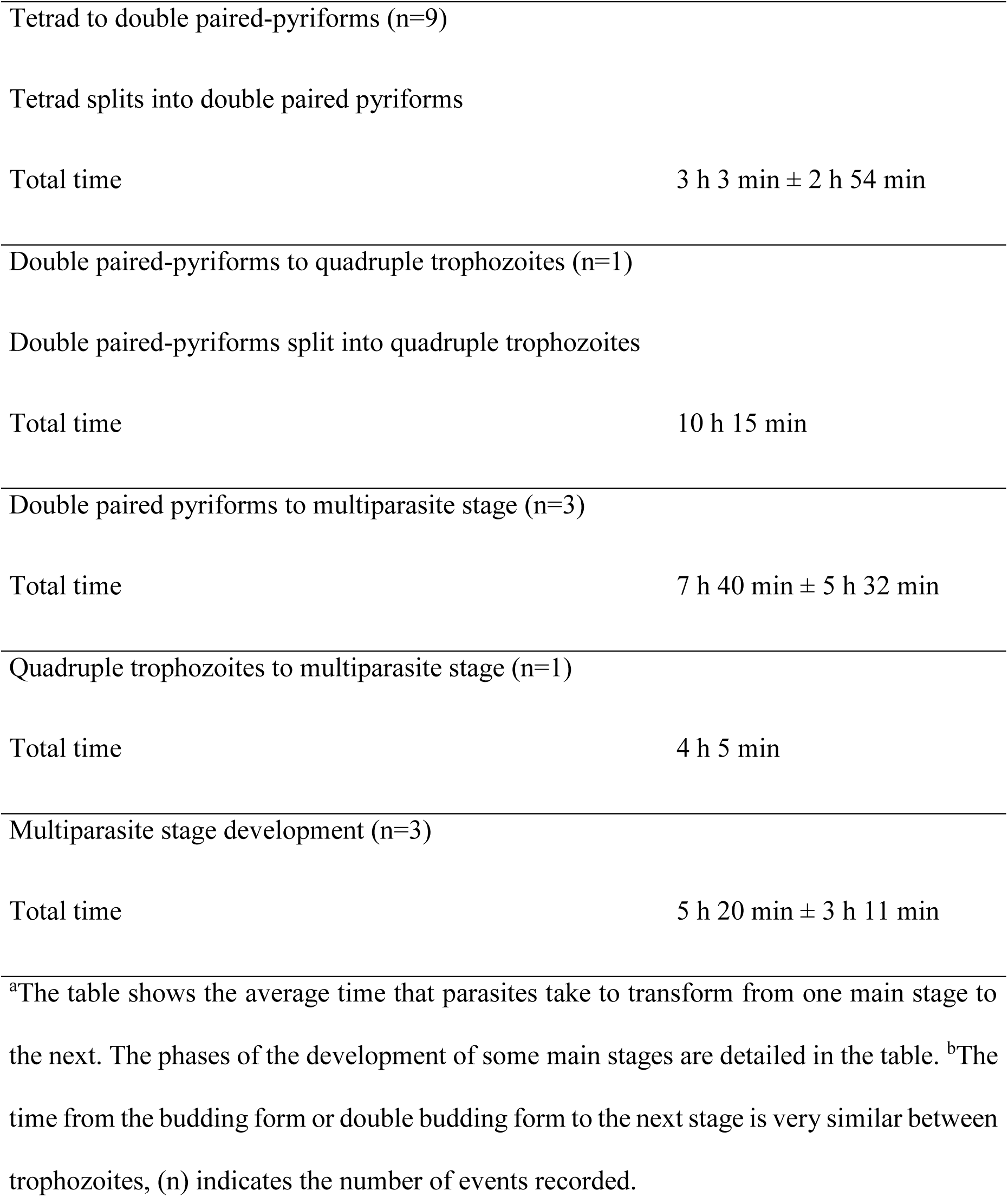
The time that intraerythrocytic parasites take to transform to the next stages.^a^

## DISCUSSION

By combining cryo-SXT and video microscopy we have obtained reconstructions and data in unprecedented detail, which significantly clarifies our understanding of the asexual cycle of *B. divergens*. The correlation between both techniques provides a new 4D vision of the cycle of native, live *B. divergens* parasites never previously visualized, thus changing our concept of parasite development in the life cycle.

The cryo-SXT tomograms revealed the main IE stages as well as unexpected forms of the parasite that were also observed and recognized by video microscopy as intermediate IE forms. These intermediate forms that interposed in a sequential order between one main stage and the next, explain the origin and development of trophozoites, pyriforms, tetrads and multiparasite stages (Movies S12-S14 at https://figshare.com/s/8ba6afd9e161899d682c).

Interestingly, the pattern of *B. divergens* population formation that we suggest here is similar to that found in blood smears of infected humans rather than infected cattle. Thus, tetrads are not typically found in cattle but are common in human RBCs and polyparasitism is also a side effect of the cycle that occurs in terminal clinical cases (6, 7, 16, 17).

The combination of video microscopy and cryo-SXT allowed the chronological ordering of a pattern of formation of the whole IE population, adopted by *B. divergens*, in the asexual cycle (Fig. 4). The proposed cycle model showed a complex morphological process where, for an individual trophozoite, there are several development options before exiting the RBC. This phenomenon occurs when parasites, instead of egressing as free merozoites to invade new RBCs, undergo several rounds of multiplication, by binary fission involving budding, within the original iRBC (Fig. 4). This phenomenon ultimately gives rise to a diverse population of multiparasite stages (Fig. 4 and Movie 14 at https://figshare.com/s/8ba6afd9e161899d682c). Our results definitely indicated that multiparasite stages initially originated from a single trophozoite, confirming that the polyparasitism phenomenon is due to continuous rounds of multiplication (17, 18) rather than multiple infections of the same RBC.

In the first phase of this process, the single trophozoite develops into a paired-pyriform rather than undergoing a duplicate binary fission event to develop into a tetrad as previously suggested (14). The new paired-pyriform precedes all the next stages that may occur in the second phase of the asexual cycle, the resulting paired-pyriform can egress from the RBC (6) or remains within the cell to develop into a tetrad or a double trophozoite. Interestingly, the tetrad exclusively derives from the paired pyriform stage. The tetrad development occurs when the paired pyriform, for unknown reasons, does not complete the fission process to separate in two trophozoites. As a consequence, the two pear-forms that encompass the paired pyriform remain attached to each other while both undergo a simultaneous but independent multiplication round involving budding. Each pear-form yield two daughter cells, four attached cells in total that form a tetrad. This could be a common cell biological feature of the *Babesia* spp. that, like *B. divergens*, are capable of producing tetrads (19, 20). Interestingly, attached or unattached, *B. divergens* parasites can just produce two daughter cells per parasite and per multiplication round.

The resulting tetrad can egress or become double paired-pyriforms within the RBC rather than develop to quadruple trophozoites. Double trophozoites, contrary to other models (7), do not leave the cell but develop into double paired-pyriforms. The latter may exit or remain within the original RBC and develop into quadruple trophozoites, a stage that does not leave the cell as well (6), but develops into a multiparasite stage. Moreover, double paired-pyriforms undergo a novel pathway consisting of sequential rounds of multiplication to yield multiparasite stages without developing first into intermediate quadruple trophozoites. Finally, multiparasite stages egress from the host cell and the resulting free merozoites invade new RBCs (Fig. 5).

**Fig. 5.**
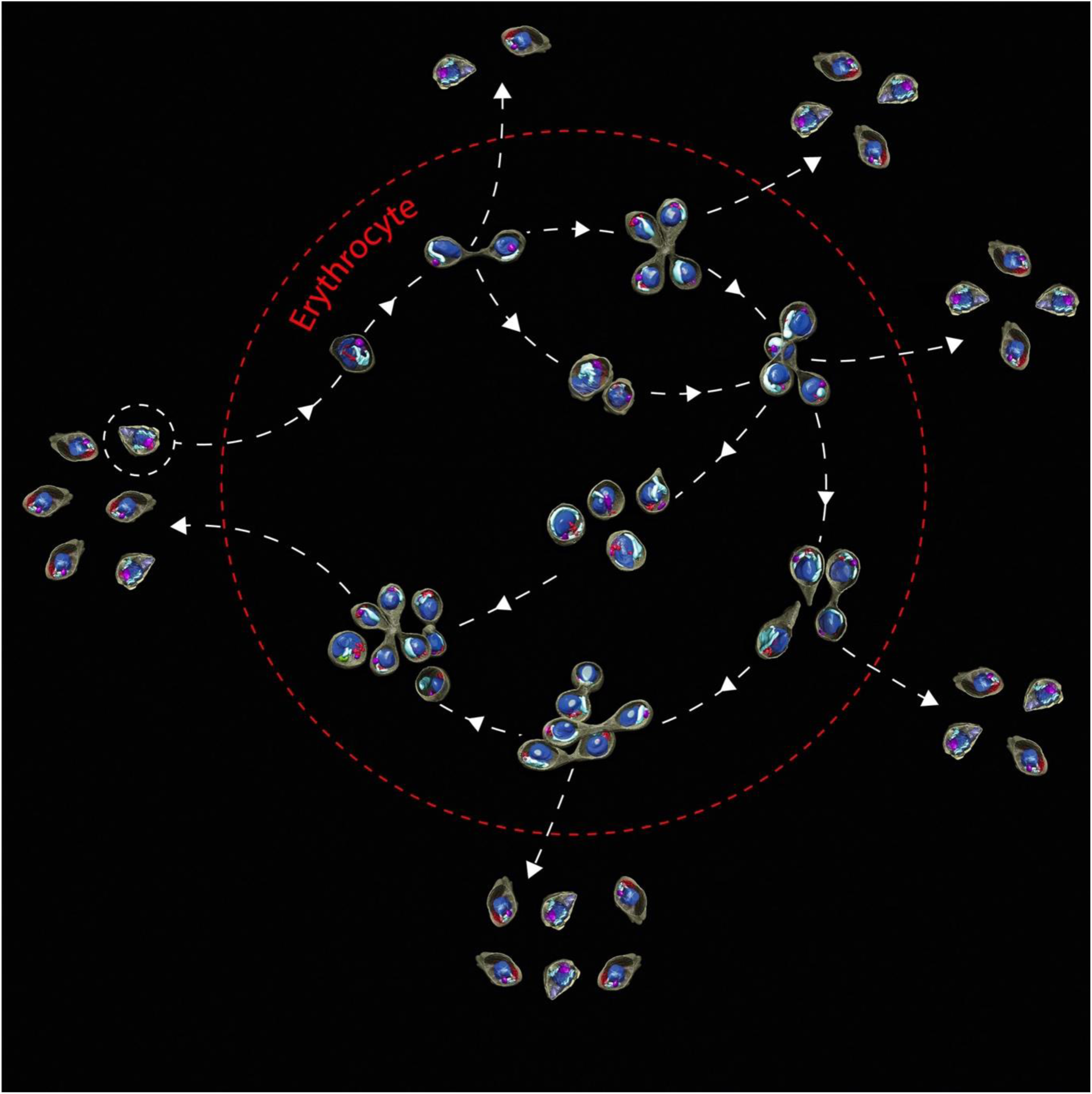
Simplified model of the *B. divergens* asexual cycle: from the single trophozoite to the multiparasite stage. The virtual model shows the transformation that a single trophozoite undergoes to become a multiparasite stage within one human RBC (red dotted line). This is possible through several rounds of multiplication by binary fission involving budding within the same host cell. This process starts with the invasion by a free merozoite (surrounded by a white dotted line) and its transformation into a single trophozoite inside the RBC. The single trophozoite develops into a paired-pyriform. This new stage develops into double trophozoites or tetrads that precede double paired-pyriforms. This last one may develop into quadruple trophozoites and/or multiparasite stages. Quadruple trophozoites can also develop into multiparasite stages. Importantly, paired-pyriforms, double paired-pyriforms, tetrads and multiparasite stages, instead of developing within the RBC, can exit the host cell as free merozoites capable of invading new RBCs resulting in a rise in parasitaemia.

Notably, the asexual cycle is an asynchronous process, and replication of two or more parasites within the same RBC does not necessarily occur simultaneously. Except for the paired pyriforms that develop into tetrads, most of the parasites multiply at different times in presence of other parasites within the same host cell. These phenomena, far from occurring as random events, follow coordinated paths, tightly controlled by the ratios of invasion, development, and egression based on environmental conditions (7, 21). Thus, under adverse conditions, or in the presence of high parasitaemias, polyparasitism increases because multiparasite stages are arrested within the host cells until favorable environmental conditions are restored (7, 21).

The main stages grow slowly and multiply, by binary fission involving budding, within the human RBC for several hours in order to achieve the next stage. The main IE forms resulting are highly active and with a great capacity for deformation and transformation to intermediate forms during the process. In this context, the integration of cryo-SXT and video microscopy data also provided a comprehensive and dynamic view of the binary fission and budding process complementary to the static view provided previously by TEM (9, 10). Indeed, it was possible to identify at least two key events to explain how fission occurs: i) the formation of a local constriction and its transformation into a protuberant knob and ultimately into a narrow waist and ii) the longitudinal stresses and the transverse rupture of the waist. While constriction formation is crucial to successful distribution of the cytoplasm to sister cells, narrow waist formation is key to rupture. Thus, a longitudinal tension force, exerted by the paired-pyriform (6), probably increases longitudinal stresses in the waist to break this structure yielding two identical pear-shaped cells concluding the fission process (Movie S14 at https://figshare.com/s/8ba6afd9e161899d682c). Fission has been recognized as a rapid process in such superior asexual organisms as freshwater planarians, which generate the forces necessary for division using only their own musculature and substrate traction (22). However, the fission process remains poorly understood in the majority of organisms because of the difficult of capturing it in real-time (22). Taking into account that *B. divergens* encodes actin, actin-polymerization and depolymerization proteins and myosin A and B (23), it is possible that cytoskeleton components, as actin-myosin motors, play a role in parasite fission.

Apart from a detailed description of the 3D morphology and kinetics of *B. divergens*, our results provide, at the nanoscale, a cell cartography associated with cytological features and profound morphological changes undergone by *B. divergens*. These include the finding of trophozoites capable of internalizing host RBC hemoglobin by the formation of a local invagination of the parasite membrane and cytoplasm, resulting in a hemoglobin-containing vesicle within the parasite. This potential avidity for hemoglobin, apparently exclusively in round trophozoites, could presumably result in later hemoglobin catabolism and production of nutrients to support *B. divergens* growth and replication during the lifecycle. While this is a well-known strategy used by *Plasmodium falciparum*, it is poorly explored in *Babesia* spp. (23, 24). Recently, relevant orthologs of the *P. falciparum* papain proteases, involved in hemoglobin digestion, were identified in the *B. divergens* genome (23) shedding more light on the role of hemoglobin during the parasite lifecycle.

After this stationary hemoglobin-phase, trophozoites apparently do not egress from the host cell but grow and multiply by binary fission in order to provide pear-shaped parasites (Fig. 5). Then, these resulting paired-pyriforms, perpetuate the cycle using two different strategies: i) leaving the cell as free merozoites in order to invade new RBCs (6), or ii) undergoing new rounds of multiplication in order to yield new IE trophozoites and pear-shaped parasites (Fig. 5, Movies S13 and S14 at https://figshare.com/s/8ba6afd9e161899d682c).

During this dynamic cyclic process, *B. divergens* stages produce sub-micron vesicles observable within both the parasites and the iRBC cytoplasm (but not seen in uRBCs), together with long membranous extensions connecting the IE parasite to the RBC plasma membrane. The presence and features of both membranous structures suggests the establishment of parasite/host cell interactions and the interchange of parasite/host cell products in a differ manner than the system of connected vesicles, used exclusively by *B. microti*– and *B. duncani*– iRBCs, for parasite antigen export (25).

Thus, both round trophozoites and pear-shaped forms are highly active and interacting parasites, and have different but complementary roles. While, trophozoites probably ensure the first nutrients by capturing hemoglobin, pear-shaped parasites seem to be the first step in perpetuating the cycle.

Undoubtedly, the life cycle of *B. divergens* requires precise strategies to ensure efficient propagation. Imaging tools showed a complex morphological presentation of IE parasites and provide a better understanding of the role that *B. divergens* performs inside its host cell.

Further exploration of the whole *Babesia* life cycle, which spans two hosts—a tick vector and a vertebrate— will be crucial to improve our knowledge of the basic biology, morphology, and host–pathogen interactions of this parasite, and to facilitate the parasite diagnosis and to provide better strategies for control.

## MATERIALS AND METHODS

### Ethics statement

Human A+ blood from healthy donors was used to maintain cultures of *B. divergens*. The blood and protocol were approved for use by the Blood Transfusion Center, Madrid, Spain. Donors provided informed written consent for use of their blood for research purposes.

### Parasite propagation

*B. divergens* asynchronous cultures (Bd Rouen 1987 strain) were maintained *in vitro* in human A+ RBCs at 5% hematocrit (9). Infected RBCs were stained with Giemsa and examined with a Primo Star microscope (Zeiss, Germany) at 100X magnification.

### Cryo-epifluorescence microscopy

Cultures of *B. divergens* at 30% parasitaemia were stained with MitoTracker Red FM mitochondrial stain (Thermo Fisher Scientific, OR, USA) at a final concentration of 500 nM and following the manufacturer’s instructions. Then, 10^5^ fluorescence-stained cells were deposited on the surface of Au-G200F1 finder grids coated with holey carbon (R 2/2; Quantifoil) and functionalized with Poly-L-Lysine (Merck, Germany) and fiducial gold markers of 100 nm (BBI Solutions, UK) used for tomographic alignment purposes. To conserve the cellular structures and membrane arrangements in close-to-native conditions, cells attached to the grids were cryo-fixed by plunge freezing in liquid ethane using a Leica EM CPC plunge freezer (Leica Microsystems, Germany). Vitrified grids were transferred in liquid nitrogen to the cryo-correlative cooling stage (CMS196 stage, Linkam Scientific Instruments, UK) to hold samples at a stable −190 °C during analysis. The cryo-stage was inserted into an AxioScope A1 (Carl Zeiss, Germany) epifluorescence microscope with an N-Achroplan 10×/0.25 Ph1 objective and imaged with a CCD AxioCam ICm1 (Carl Zeiss). Cryo-fluorescence correlative microscopy was used to pre-select vitrified samples and map cell coordinates. Selected samples were then transferred to ALBA synchrotrons at liquid nitrogen temperature.

### Cryo-Soft X-ray tomography

Holey carbon-coated (R 2/2; Quantifoil) Au-G200F1 grids were analyzed in cryo-conditions by MISTRAL microscope at ALBA synchrotron, Barcelona, Spain (13). RBCs infected with red fluorescence *B. divergens* parasites were visualized on-line with a transmitted visible light and epi-fluorescence microscope integrated within the Mistral Soft X-Ray Microscope to re-map cell coordinates and select the cryo-SXT acquisition areas. X-ray projection mosaics were acquired to evaluate sample vitrification and thickness. Tilt series were acquired from −65° to 65° at 1° intervals, using a 25 nm zone plate objective lenses. Exposure time was 1-2 s, depending on sample thickness, and the effective pixel size 10 nm. Additionally, some samples were mounted in AutoGrid supports (FEI) and imaged following a dual axis acquisition scheme. Most single axis acquisition tomograms were done following a XTEND acquisition scheme (11). We imaged 218 acquisition areas as follows: 26 single axis tilt series, 42 dual axis tilt series and 150 XTEND tilt series.

Tilt series were normalized to the flatfield, deconvolved by the measured apparent transfer function of the microscope (26) using python and MATLAB scripts and aligned with IMOD (27). XTEND data series were processed as described (28) using python scripts. Tomographic reconstructions were performed using TOMO3D software, 30 iterations of simultaneous iterative reconstruction technique (SIRT) algorithm (29) and edge enhanced using TOMOEED (30). Segmentation of volumes was carried out with SuRVoS (31), and volumes were represented with Chimera (32) and ImageJ (33).

### Staining *B. divergens* culture parasites with MitoTracker green and subsequent treatment with Concanavalin A

*B. divergens* cultures of at 25-28% parasitaemia were stained with MitoTracker Green FM mitochondrial stain (Thermo Fisher Scientific) at a final concentration of 500 nM (6). Culture samples were placed in 6-well cell-culture plates and maintained at 37 °C in a humidified atmosphere of 5% CO2 until use. Then, wells of a glass-bottom 96-Well Black, No. 1.5 Coverslip, 5 mm Glass Diameter, Uncoated (MatTek, MA, USA) were treated with 50 *μ*l of Concanavalin A (Sigma-Aldrich Corporation, St Louis, MO), at 0.5 mg/ml, for 10 minutes at room temperature and washed twice with 200 *μ*l of PBS 1x. Simultaneously, the RBCs infected with green fluorescence *B. divergens* parasites were also washed in PBS 1x. Cells (5×10^5^-1×10^6^ per well) were placed in the wells and stuck for 5 minutes at room temperature. Then, non-bound cells were removed and bound cells were washed twice with 200 *μ*l of PBS 1x. Finally, PBS 1x was replaced by 200 *μ*l of complete medium to maintain the culture during the video microscopy assays.

### Long-term time-lapse recording and video processing

Time-lapse video was conducted using a Leica TCS SP5 confocal laser microscope (Leica Microsystems) equipped with epifluorescence microscopy (Leica DMI 6000B microscope) and incubation systems to control temperature, humidity and CO2 conditions. To avoid loss of focus during the video recording, a 96-well plate, containing RBCs infected with green fluorescence *B. divergens* parasites, was placed under the confocal microscope with 63x oil objective lens and incubated in a 5% CO2 environment at 37 °C for 1 hour.

Time-lapse images of iRBCs were then recorded at one frame per 5 minutes interval using the following parameters: 488 nm laser line and laser level of 10%, a speed of 700 Hz, a 2.25 AU pinhole aperture, a zoom of 2x, 2.5x or 3x and bright field imaging under the same environmental conditions. Frames were captured for 18-21 hours in a single z-section. The videos generated by the LAS AF software were processed with ImageJ and Fiji software (33, 34).

### Transmission electron microscopy (TEM)

For TEM ultrastructural analysis, *B. divergens in vitro* cultures were stuck to microscope cover glasses (12 mm) using poly-L-lysine (Merck). Briefly, samples were fixed in 2.5% glutaraldehyde and 2% paraformaldehyde in 0.1 M Na2HPO4, pH 7.4; post-fixed with 1% osmium tetroxide and 1% uranyl acetate, dehydrated in increasing concentrations of ethanol, infiltrated using increasing concentrations of epoxy-resin and polymerized at 60 °C for 48 h. Serial ultra- and semi-thin sections (70–150 nm) were obtained with a Leica EM UC6 ultramicrotome and harvested following standard procedures (8). Images were registered on a FEI Ceta camera with a Tecnai 12 FEI microscope operated at 120 kV.

### Statistical analysis

Mean and standard deviations (SD) were calculated using Excel 2010 (Microsoft, Redmond, WA, USA) and results were expressed as average ± SD.

### Data availability

Supplemental materials (Movies S1-S14) are available at Figshare: https://figshare.com/s/8ba6afd9e161899d682c

## ACKNOWLEDGMENTS

We thank Centro de Transfusiones de la Comunidad de Madrid which provided the human A+ blood from healthy volunteer donors and V. Lavilla for designing the animation movies. This work was funded by grants from Ministerio de Economía y Competitividad from Spain (AGL2010-21774, AGL2014-56193-R to EM and LMG, BFU2013-43149-R to DL). Cryo-SXT experiments were funded by ALBA synchrotron from Barcelona, Spain (Proposals 2016021614 and 2017022084) and performed at MISTRAL beamline at ALBA Synchrotron with the collaboration of ALBA staff. E. Sevilla was awarded a research fellowship from Plan Estatal de Investigación Científica y Técnica y de Innovación.

## AUTHORSHIP CONTRIBUTIONS

J.J.C., L.M.G., E.P., F.J.C., D.L. and E.M. designed research; J.J.C., E.S., M.C.T., L.M.G., J.G., A.J.P.B., J.L.C., E.P., F.J.C., D.L. and E.M. conducted experiments and/or analysis; J.J.C., D.L. and E.M. wrote the paper; J.J.C. and E.S. made figures; J.J.C., E.S., M.C.T., L.M.G., A.J.P.B., J.G., J.L.C., E.P., F.J.C., D.L. and E.M. conducted review and editing; L.M.G., D.L., E.P. and E.M. provided funding acquisition, project administration, and resources.

## SUPPORTING INFORMATION CAPTIONS

**Fig. S1. Cryo-SXT tomography of the *B. divergens* free merozoite**. (a-c) Different Cryo-STX sections from a tomogram of a free merozoite. Arrow heads point to putative structures and secreted organelles, related to invasion and egress of parasites that were classified according to their differences in dimension and disposition within the parasite. (a) Inner membrane complex. (b) Micronemes (red arrow head) and rhoptries (white arrow head). c) Dense granules (white arrow heads). The scale bar is 500 nm.

**Fig. S2. Cryo-SXT and TEM analysis of possible hemoglobin-containing vesicles**. (a-b) Cryo-STX sections of RBCs infected with *B. divergens* round trophozoites containing dense structures (white head arrows). These structures exhibit similar X-ray absorption coefficients to the hemoglobin from the RBC cytoplasm. (c-e) Serial sectioning TEM images of the endocytic uptake of the hemoglobin (white head arrows) by a trophozoite. (f-h) Serial sectioning TEM images of the resulting hemoglobin-filled vesicle within the trophozoite (white head arrows). The scale bars of a and b are 2 µm. The scale bars of c-h are 500 nm.

**Fig. S3. Cryo-SXT and TEM analysis of sub-micron vesicles present in the cytoplasm of *B. divergens* human iRBCs**. (a-d) Cryo-SXT tomogram sections of erythrocytes infected with different *B. divergens* stages. (a) Trophozoite. (b) Multiparasite stage. (c) Trophozoite. (d) Paired-pyriform. (e-h) TEM images showing RBCs infected with different *B. divergens* forms. The white arrow heads point to sub-micron vesicles ranging from 120 to 400 nm within the cytoplasm of the iRBCs. The scale bars of a-d are 2 µm. The scale bars of e-h are 500 nm.

**Fig. S4. Cryo-SXT and TEM analysis of long membrane structures. (a-c)** Cryo-SXT tomogram sections of RBCs infected with different *B. divergens* stages. (a) Trophozoite. (b and c) Two different cryo-SXT tomogram sections of a paired-pyriform. (d-f) TEM images showing *B. divergens* iRBCs. The white arrow heads from a-c panels point to a long membrane structure that makes contact with both the parasite and RBC plasma membranes. The white arrow heads from panels d-f point to similar long membranous structures to those indicated in a-c panels. The scale bars of a-c are 2 µm. The scale bars of d-f are 1 µm.

**Fig. S5. Cryo-SXT of *Babesia divergens* paired-pyriforms. (**a-c) Panels show cryo-STX tomogram sections of paired-pyriforms within RBCs. (d, e) Panels show TEM sections of paired-pyriforms within the RBC. Subcellular structures are visible within a protuberant knob (white arrows) that is formed during the paired-pyriform development. (c) The knob ultimately develops into a narrow waist that connects both pear-shaped parasites. The scale bars of a-e are 1 □m.

**Fig. S6. Developing process of *B. divergens* double trophozoite and double paired-pyriform stages**. The figure shows the development of double trophozoites from the paired-pyriform (panel a) and the development of double paired-pyriforms (panel b). Transient forms that intersperse in a chronological order, between the main stages, are represented in the figure. The top row shows time lapse image sequences, captured by video microscopy, of parasites stained with MitoTracker (green fluorescence) transforming within the RBCs. The serial diagrams (middle row) and Giemsa-stained parasites (bottom row) recreate the development process. (a) A paired-pyriform separates transversely at 7 h yielding two identical pear-shaped sister cells. The pear-shaped cells adopt different forms until reaching the characteristic round shape of trophozoites at 10 h and 55 min. The whole development process results in double trophozoites. (b) Two pear-shaped cells transform into trophozoites at 5 h and 5 min. The resulting trophozoites undergo a non-simultaneous multiplication by binary fission involving budding. The first trophozoite (outlined in red) reaches the budding form at 10h and 30 min and ultimately develops into a paired-pyriform at 12 h and 45 min. The second trophozoite (outlined in white) reaches the budding form 2 hours and 15 min later than the first one (at 12 h and 45 min) and ultimately develops into a paired-pyriform at 15 h. The whole development process results in double paired-pyriforms. Time-lapse imaging was captured every 5 min. The time lapse between each frame is indicated in hours and minutes. The scale bar is 5 µm (see also Movies S2 and S3 at https://figshare.com/s/8ba6afd9e161899d682c)

**Fig. S7. Developing process of the double paired-pyriform stage**

The figure shows the development of the double paired-pyriform stage and the transient forms that appear in a chronological order. The top row shows time lapse image sequences, captured by video microscopy, of parasites stained with MitoTracker (green fluorescence) transforming within the RBC. The serial diagrams (middle row) and Giemsa stained parasites (bottom row) recreate the development process. The first panel (00:00 h) shows a RBC infected with a paired-pyriform and a trophozoite. While the paired-pyriform (originally from a trophozoite) remains in a stationary phase for 4 h and 40 min, the trophozoite adopts an amoeboid shape until it reaches the budding form at 2 h. In a second phase of development, the budding form becomes elongated, meanwhile a transverse constriction is formed around the body. The transverse constriction changes to a protuberant knob at 2 h and 30 min. The knob is progressively transformed into a narrow waist at 3 h and 25 min. The main body is constricted, through the waist, forming two attached pear-shaped sister cells at 4 h and 40 min. The development process results in double paired-pyriforms. Time-lapse imaging was captured every 5 min. The time lapse between each frame is indicated in hours and minutes. The scale bars is 5 µm (see also Movie S4 at https://figshare.com/s/8ba6afd9e161899d682c).

**Fig. S8. Paired-pyriforms developing into tetrads and double paired-pyriform stages**. The figure shows the development of paired-pyriforms into tetrads and into double paired-pyriforms. Transient forms appear in a chronological order between main stages. Top rows show time lapse image sequences, captured by video microscopy, of parasites stained with MitoTracker (green fluorescence) transforming within the RBC. Parasite forms are outlined in white to facilitate the monitoring of the sequential events. The serial diagrams (middle row) and Giemsa-stained parasites (bottom row) recreate the development process. (a) Panels show an RBC infected with a paired-pyriform developing into a tetrad. The two sister pear-shaped forms that comprise the paired-pyriform remain attached and simultaneously adopt amoeboid forms until yielding a double budding form at 3 h and 5 min. In a second phase of development, the double budding form undergoes a morphological transformation and develops into a tetrad at 4 h and 10 min. The resulting tetrad is formed by four attached pear-shaped sister cells. (b) Panels show an RBC infected with a paired-pyriform developing into a tetrad that ultimately develops into double paired-pyriforms. The paired-pyriform remain attached and simultaneously adopt amoeboid forms until yield a double budding form at 7 h and 5 min and ultimately reaches a tetrad at 10 h and 10 min. Notably, the tetrad separates in the middle yielding a double paired-pyriform stage at 10 h and 25 min. Time-lapse imaging was captured every 5 min. The time lapse between each frame is indicated in hours and minutes. The scale bars is 5 µm (see also Movies S5 and S6 at https://figshare.com/s/8ba6afd9e161899d682c).

**Fig. S9. Dynamic activity of tetrad and double paired-pyriform stages**. The figure shows the development of a tetrad into a double paired-pyriform stage and the kinetics of this last stage within the RBC. Top rows show time lapse image sequences, captured by video microscopy, of parasites stained with MitoTracker (green fluorescence) transforming within the RBC. The serial diagrams (middle row) and Giemsa stained parasites (bottom row) recreate the development process. (a) Panels show an RBC infected with a tetrad. The four sister pear-shaped forms that comprise a tetrad remain attached until they separate in pairs yielding the double paired-pyriform stage at 4 h. (b) Panels show an RBC parasitized with a double paired-pyriform stage. While one of the paired-pyriforms that comprise this stage remains in a stationary phase, the second one separates transversely along its narrowest part at 2 h and 20 min resulting into two single pear-shaped sister cells. Time-lapse imaging was captured every 5 min. The time lapse between each frame is indicated in hours and minutes. The scale bars is 5 µm (see also Movies S7 and S9 at https://figshare.com/s/8ba6afd9e161899d682c).

**Fig. S10. The development process of the multiparasite stage**. The figure shows the development of multiparasite stages and transient forms that appear in a chronological order during the process. Top rows show time lapse image sequences, captured by video microscopy, of parasites stained with MitoTracker (green fluorescence) transforming within the RBC. The serial diagrams (middle row) and Giemsa stained parasites (bottom row) recreate the development process. (a) Panels show an RBC infected with a paired-pyriform and two amoeboid trophozoites. While the paired-pyriform (outlined in blue) remains in a stationary phase for 5 h, both trophozoites develop into paired pyriforms. The trophozoite outlined in white becomes a budding form at 1h and 45 min and transforms in an early paired pyriform at 2 h and 20 min. This early paired-pyriform progresses until a mature paired-pyriform at 5 h. The second trophozoite (outlined in red) multiplies by binary fission involving budding but later at 2 h and 20 min, giving rise to a mature paired-pyriform at 5 h. The whole process results in the formation of a multiparasite stage composed by a trio of paired-pyriforms. (b) Panels show an RBC infected with an amoeboid trophozoite (outlined in blue), an early paired-pyriform (outlined in red) and a paired-pyriform (outlined in white). The continuous rounds of multiplication by binary fission involving budding that parasites undergo ultimately provoke a polyparasitism phenomenon into the RBC. The trophozoite and the early paired-pyriform become mature paired-pyriforms at 2 h and 10 min. Then, paired-pyriforms outlined in red and white separate into single pear-shaped sister cells. The two single pear-shaped sister cells (outlined in white) undergo new rounds of multiplication, but not simultaneously, yielding two new paired-pyriforms (outlined in white) at 7 h and 35 min. Time-lapse imaging was captured every 5 min. The time of each frame is indicated in hours and minutes. The scale bars is 5 µm (see also Movies S10 and S11 at https://figshare.com/s/8ba6afd9e161899d682c).

**Movie S1. From the single trophozoite to the paired pyriform**. Time-lapse video microscopy of a single trophozoite that develops into a paired-pyriform stage. Frames are captured every 5 min and the time is indicated in hours and minutes. The video is showed on two individual channels together with the merged images. The scale bars is 5 µm.

https://figshare.com/s/8ba6afd9e161899d682c

**Movie S2. The double trophozoite stage development**. Time-lapse video microscopy of a paired-pyriform that develops into a double trophozoite stage. Frames are captured every 5 min and the time is indicated in hours and minutes. The video is showed on two individual channels together with the merged images. The scale bars is 5 µm.

https://figshare.com/s/8ba6afd9e161899d682c

**Movie S3. The double paired-pyriform stage development**. Time-lapse video microscopy shows double trophozoites that develop into double paired-pyriforms. The video shows the non-simultaneous multiplication of both trophozoites. Frames are captured every 5 min and the time is indicated in hours and minutes. The video is showed on the green fluorescence channel together with the merged images. The scale bars is 5 µm.

https://figshare.com/s/8ba6afd9e161899d682c

**Movie S4. Cellular multiplication by binary fission**. Time-lapse video microscopy shows a detailed process of multiplication of a trophozoite within a parasitized RBC. Frames are captured every 5 min and the time is indicated in hours and minutes. The video is showed on two individual channels together with the merged images. The scale bars is 5 µm.

https://figshare.com/s/8ba6afd9e161899d682c

**Movie S5. The tetrad stage development**. Time-lapse video microscopy shows a paired-pyriform that develops into a tetrad. Frames are captured every 5 min and the time is indicated in hours and minutes. The video is showed on two individual channels together with the merged images. The scale bars is 5 µm.

https://figshare.com/s/8ba6afd9e161899d682c

**Movie S6. From the paired-pyriform to the double paired-pyriform stage**. Time-lapse video microscopy shows a paired-pyriform that develops into a tetrad and ultimately into a double paired-pyriform stage. Frames are captured every 5 min and the time-lapsed between each frame is indicated in hours and minutes. The video is showed on two individual channels together with the merged images. The scale bars is 5 µm.

https://figshare.com/s/8ba6afd9e161899d682c

**Movie S7. From the tetrad to the double paired-pyriforms** Time-lapse video microscopy of a tetrad that develops into a double paired-pyriforms. Frames are captured every 5 min and the time is indicated in hours and minutes. The video is showed on two individual channels together with the merged images. The scale bars is 5 µm.

https://figshare.com/s/8ba6afd9e161899d682c

**Movie S8. From the paired-pyriform to the quadruple trophozoite stage**. Time-lapse video microscopy shows double paired-pyriforms that ultimately separate into four identical sister cells that start to develop into trophozoites. Frames are captured every 5 min and the time is indicated in hours and minutes. The video is showed on two individual channels together with the merged images. The scale bars is 5 µm.

https://figshare.com/s/8ba6afd9e161899d682c

**Movie S9. The double paired-pyriforms stage separate to form single pear-shaped cells**. Time-lapse video microscopy shows two paired-pyriforms within an RBC. While one paired-pyriform remains in a stationary phase, the other one separates into two pear-shaped sister cells. Frames are captured every 5 min and the time is indicated in hours and minutes. The video is showed on two individual channels together with the merged images. The scale bars is 5 µm.

https://figshare.com/s/8ba6afd9e161899d682c

**Movie S10. The development process of the multiparasite stage**. Time-lapse video microscopy shows the continuity of the process that takes places in video 9. Video 10 shows an RBC infected with a paired-pyriform and two trophozoites. Both trophozoites multiply by binary fission, involving budding, yielding a multiparasite stage that ultimately exits the host cell. Frames are captured every 5 min and the time is indicated in hours and minutes. The video is showed on two individual channels together with the merged images. The scale bars is 5 µm.

https://figshare.com/s/8ba6afd9e161899d682c

**Movie S11. The development process of the multiparasite stage**. Time-lapse video microscopy shows a polyparasitized RBC. Some of the parasites inside the host cell multiply by binary fission, involving budding, provoking a polyparasitism phenomenon within the RBC. The multiparasite stage ultimately exits the host cell. Frames are captured every 5 min and the time is indicated in hours and minutes. The video is showed on two individual channels together with the merged images. The scale bars is 5 µm.

https://figshare.com/s/8ba6afd9e161899d682c

**Movie S12**. Schematic movie showing a trophozoite that develops into a paired-pyriform.

https://figshare.com/s/8ba6afd9e161899d682c

**Movie S13**. Schematic movie showing a paired-pyriform that develops into a tetrad.

https://figshare.com/s/8ba6afd9e161899d682c

**Movie S14**. Schematic movie showing the multiparasite stage development.

https://figshare.com/s/8ba6afd9e161899d682c

